# Cinnamaldehyde interacts with the local anesthetic pocket of the Na_V_ 1.5 channel

**DOI:** 10.64898/2026.09.13.751249

**Authors:** Julio Alvarez-Collazo, Alejandro López Requena, Ariel Talavera, Julio L. Alvarez, Karel Talavera

## Abstract

Cinnamaldehyde (CA) is extensively used as flavorant and in traditional medicine. CA is commonly used in pain research as specific agonist of TRPA1, a polymodal cation channel expressed in nociceptive primary sensory neurons. However, we previously showed that CA inhibits the L-type Ca^2+^ channel in cardiac and smooth muscle cells, raising the possibility that other ion channels can be modulated by this compound as well. Here, we investigated whether CA exhibits affects the activity on the cardiac Na_V_1.5 channel. We used the whole-cell patch-clamp technique to record Na^+^ currents in HEK293T cells expressing the human Na_V_1.5 channel, as well as in cells expressing hNa_V_1.5 channels baring mutation in the binding pocket of local anesthetics (LA). Our results show that CA exhibit LA-like actions on hNa_V_1.5 channels: CA blocks hNa_V_1.5 currents in a tonic and voltage-dependent fashion. Residues F1760 and Y1767 are important for the blockade of Na_V_1.5 by CA and a double point mutation F1760A/Y1767A abolishes the blockade of I_Na_ by CA. We conclude that CA inhibits the cardiac Na_V_1.5 channel by interacting with the LID binding site and that CA has LA-like actions.

## 1. Introduction

Cinnamaldehyde (CA), the main component (∼90 %) of cinnamon extracts, is a natural compound extensively used as a flavorant [1], as well as in traditional medicine as antimicrobial, antidiabetic and anticancer [2-4]. CA is also a TRPA1 cation channel agonist [5] commonly used in pain research. However, we have previously shown that CA inhibits the bacterial voltage-gated Na^+^ channel NaChBac [6]. This raises the possibility that this compound inhibits also other voltage-gated Na^+^ channels. Curiously, it has been shown that LID activates TRPA1 channels with an EC_50_ of ∼ 5 mmol·L^-1^ for current activation in HEK293 cells [7]. The residue F1760 in the main cardiac Na^+^ channel isoform (Na_V_1.5), has been identified as the main structural determinants for the high-affinity block of Na_V_ channels by LA [8-10]. It has been proposed that LA-like drugs bind to the F1760 residue via a strong electrostatic cation–π interaction [11]. LA bound to this region may prevent the current flow through Na^+^ channels by introducing a positive charge, blocking Na^+^ movement in an electrostatic way [12, 13]. Although the residue F1760 stands out for playing a key role in the binding of LA-like compounds some studies reveal that point mutations in the Y1767 residue could also participate in modulating the effect of LA-like drugs on Na^+^ channels [14, 15]. However, the possible interactions of CA with other residues in the LA pocket as well as its LA-like actions on the Na_V_1.5 channel have not been investigated. Given the structural and functional similarities between CA and LID, the aim of the present work was to study the possible interactions of CA with the residues lining the LA pocket in the human Na_V_1.5 (hNa_V_1.5) channel. Our results show that CA exhibits a LA-like blocking action on hNa_V_1.5 channels and that the residues F1760 and Y1767 play an important role in this action.

## 2. Results

### 2.1. Blockade of WT human Na_V_1.5 channels by cinnamaldehyde

We first studied the effects of CA on I_Na_ in HEK293T cells expressing the WT hNa_V_1.5 channel. In control condition, peak inward I_Na_ density at -20 mV was -112 ± 18 pA·pF^-1^ with a mean time to peak of 1.07 ± 0.05 ms (n = 39). Under continuous voltage-clamping (0.25 Hz), CA induced a concentration-dependent decrease of peak I_Na_ (Figure 1A). Blockade occurred in two phases, a fast one-pulse block that accounts for 76.9 ± 2.9 % (n = 13) of total blockade at CA 3 mmol·L^-1^ followed by a slower block that reached a steady state in ≈ 20 s. The washout of CA was in an “off” fashion. Experimental data using different CA concentrations were fitted by a Hill function giving an IC_50_ for the steady-state blockade of peak I_Na_ of 1.04 ± 0.08 mmol·L^-1^ (Hill number, H = 1.7 ± 0.22; Figure 1B). Time to peak I_Na_ was not affected by CA, even at the highest concentration applied (3 mmol·L^-1^; 1.03 ± 0.08 ms, n = 13).

**Figure 1.**
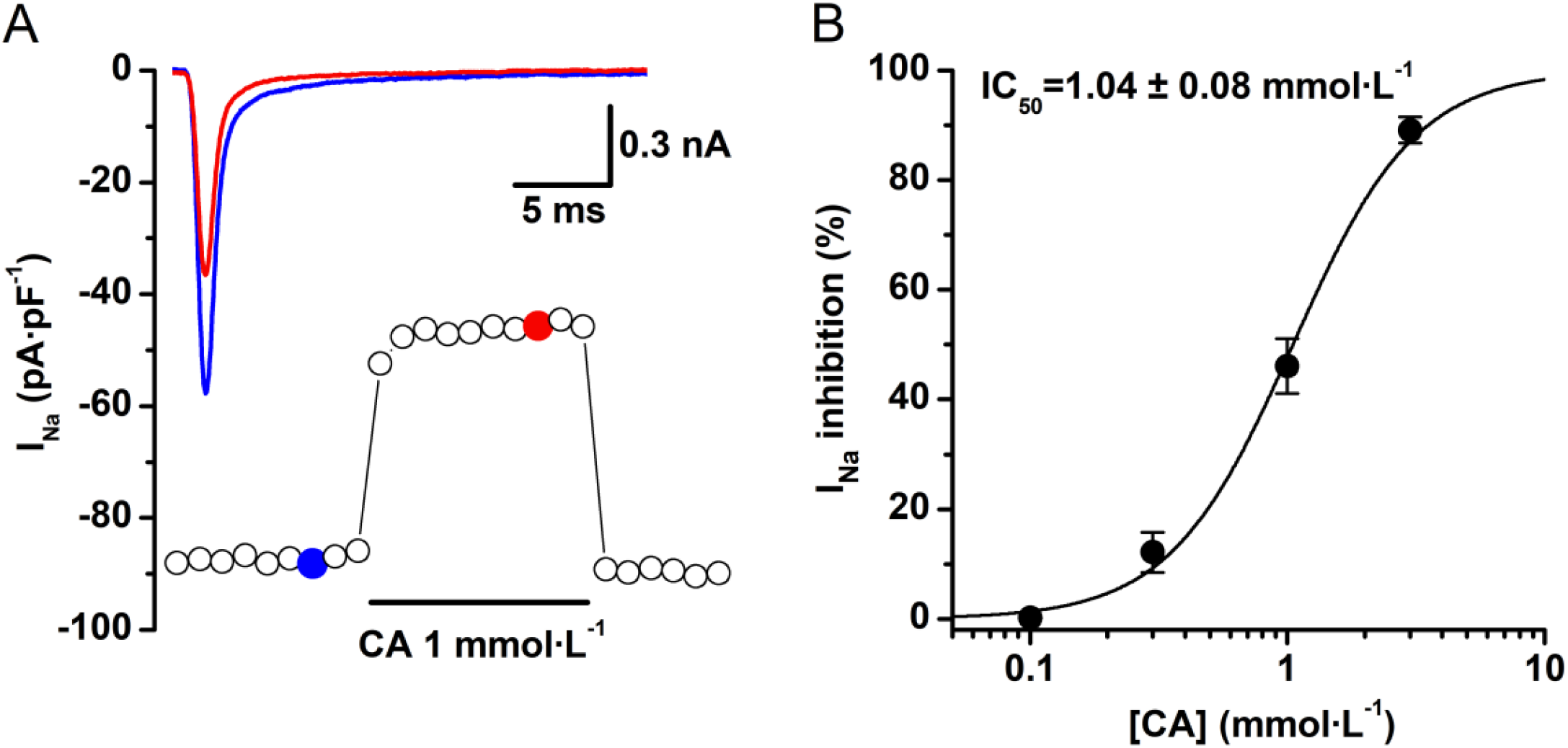
Cinnamaldehyde blocks I_Na_ in HEK293T transiently expressing the hNa_V_1.5 channel. A.-Time course of the CA effect on the I_Na_ from a HEK293T cell expressing the WT hNa_V_1.5. CA 1 mmol·L^-1^ was applied as indicated. Representative current traces in each condition are shown in the inset and correspond with the colored data points: blue, control; red, CA. For clarity only the first 25 ms of the current traces are shown. B.-Concentration - effect relationship for the inhibitory action of CA on hNa_V_1.5 current. The dots represent the mean ± s.e.m. inhibition percentage during the application of different concentrations of CA (n = 13). The line represents the Hill fit of the data and IC_50_ value is shown on the inset.

We studied in detail tonic (TB) and use-dependent (UDB) blockades of hNa_V_1.5 by CA. In control conditions after a sudden increase in the frequency of stimulation there were no significant variations in I_Na_ amplitude but, when CA (1 mmol·L^-1^) was perfused, a huge TB followed by a small UDB could be observed (Figure 2A). TB amounted to 92.0 ± 3.3 % of the corresponding total block which, at this concentration, was near 50 % of control I_Na_ (n = 6). At a stimulation rate of 1 Hz, the additional UDB was only 8.0 ± 3.3 % (Figure 2B), i.e. at the highest frequency of stimulation, total block of I_Na_ by CA was almost no further enhanced (Figure 2C).

**Figure 2.**
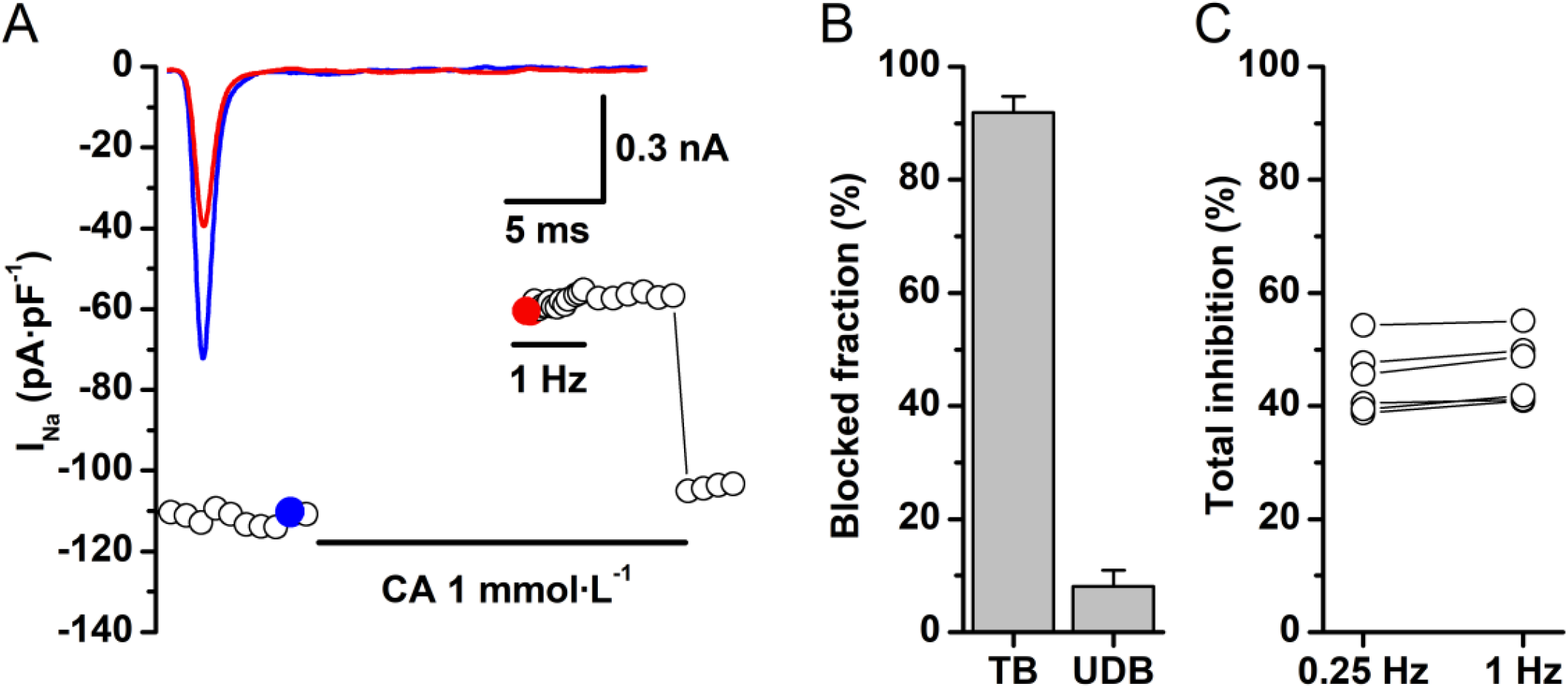
Characteristics of the block of I_Na_ by cinnamaldehyde. A.-The cells were clamped at -100 mV and test pulses to -20 mV were applied at 0.25 Hz. Stimulation was stopped and CA (1 mmol·L^-1^) was perfused. After one-minute rest period, the same stimulation protocol was resumed but at 1 Hz and then back at 0.25 Hz. CA was constantly perfused during and after the rest. A huge tonic (resting-state) block of I_Na_ occurred when stimulation was resumed at 1 Hz. No recovery of I_Na_ amplitude was observed when stimulation rate was decreased to 0.25 Hz indicating that the inhibition of I_Na_ by CA was mostly tonic with a small use-dependent blockade. The insets show the corresponding current tracings in control (blue) and in the presence of CA 1 mmol·L^-1^ (red); for clarity only the first 25 ms of the current traces are shown. B.-Bar graph showing the blockade fraction of tonic and additional use-dependent block with respect to total block at CA 1 mmol·L^-1^. Data is presented as mean ± s.e.m. of n = 6. C.-Dot and line graph showing that CA inhibition of I_Na_ is frequency-independent. The percent of total block was almost the same at frequencies of 0.25 Hz and 1 Hz (n = 6).

Blockade of I_Na_ by CA was voltage-dependent; decreasing the HP to -80 mV, increased inhibition of I_Na_ by CA 1 mmol·L^-1^ to 64.3 ± 1.2 % (P < 0.05; n = 6; paired t-test). To further characterize the voltage-dependent effects of CA, I-V relationships, availability and activation curves were analyzed under control condition and in the presence of CA 1 mmol·L^-1^ (Figure 3A and B). In control condition the voltage of half-maximal inactivation (V_inac_) and the inactivation slope factor (s_inac_) were -72.4 ± 0.4 mV and 6.1 ± 0.4 mV, respectively. The estimated voltage of half-maximal activation (V_act_) and the activation slope factor (s_act_) were -38.3 ± 1.1 mV and 4.4 ± 0.9 mV, respectively. CA significantly shifted (P < 0.05) the V_inac_ to -84.4 ± 0.5 mV with minor changes in the slope factor (6.2 ± 0.4 mV; Figure 3B). CA did not modify V_act_ or s_act_ (-37.4 ± 1.7 mV; 4.7 ± 1.5 mV).

**Figure 3.**
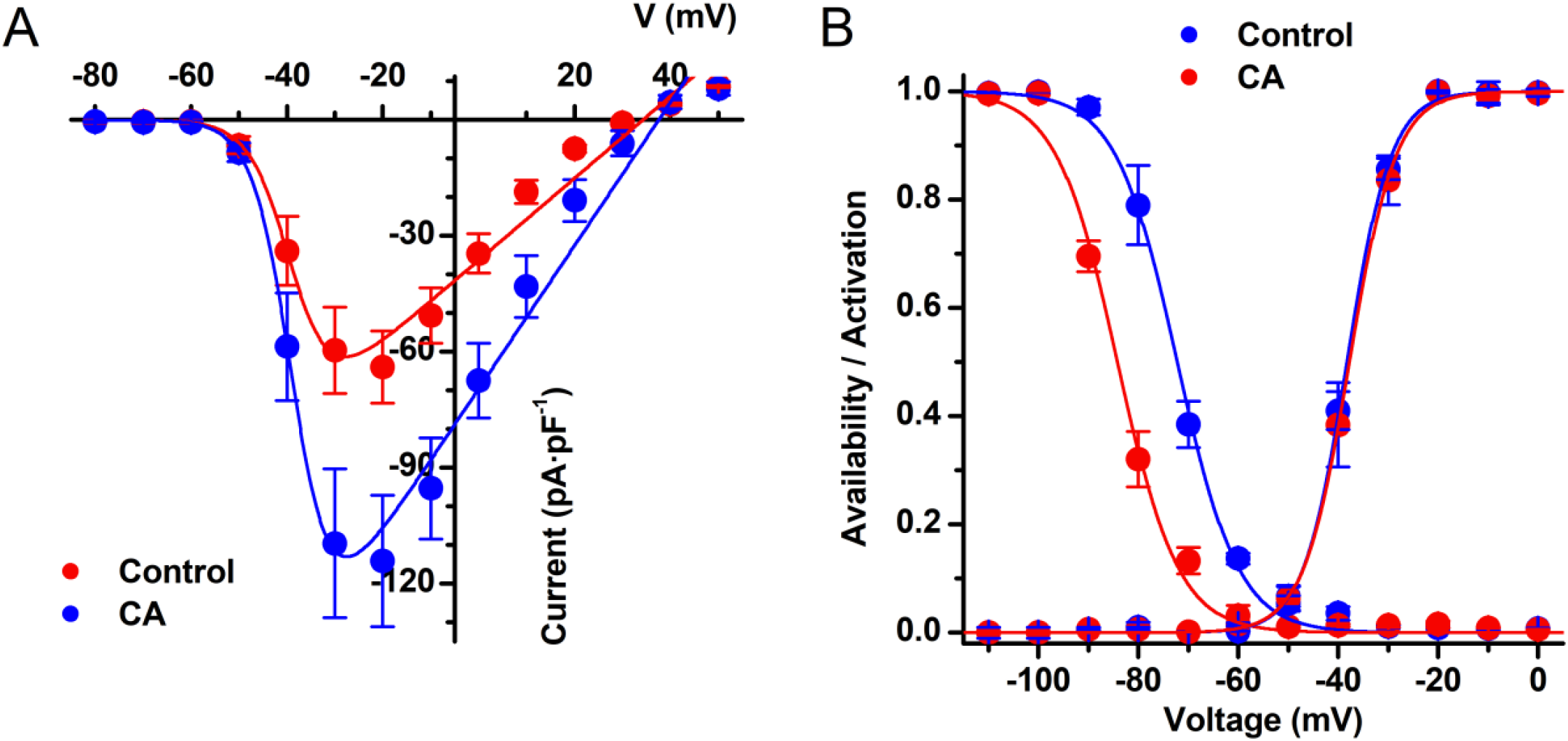
Cinnamaldehyde shifts the voltage-dependent inactivation of hNa_V_1.5. A.- Current-to-voltage relationships of I_Na_ in WT under control condition (blue) and in the presence of CA 1 mmol·L^-1^ (red). The dots represent the mean ± s.e.m. CA significantly decreased the current density compared to control in the voltage range from -40 to +20 mV (P < 0.05, n = 8; paired t-test). B.- Availability and activation curves in control (blue) and in the presence of 1 mmol·L^-1^ CA (red). The dots represent the mean ± s.e.m. of n = 8 cells. CA significantly shifted the availability curve with respect to control (P < 0.05, paired t-test).

### 2.2. Docking of cinnamaldehyde

In our docking model we found that in 9 out of 10 simulations CA was bound at distances (4.4 Å) close to the residue F1760, located in the S6 segment of the DIV of the channel, identified as the main structural determinant of the high-affinity block by lidocaine [8, 9] (Figure 4). However, the docking model also suggests that the residues F934 (at 3.7 Å), F1465 (at 3.6 Å) and Y1767 (at 3.6 Å) could be also important in determining CA blockade of the Na_V_1.5 channel.

**Figure 4.**
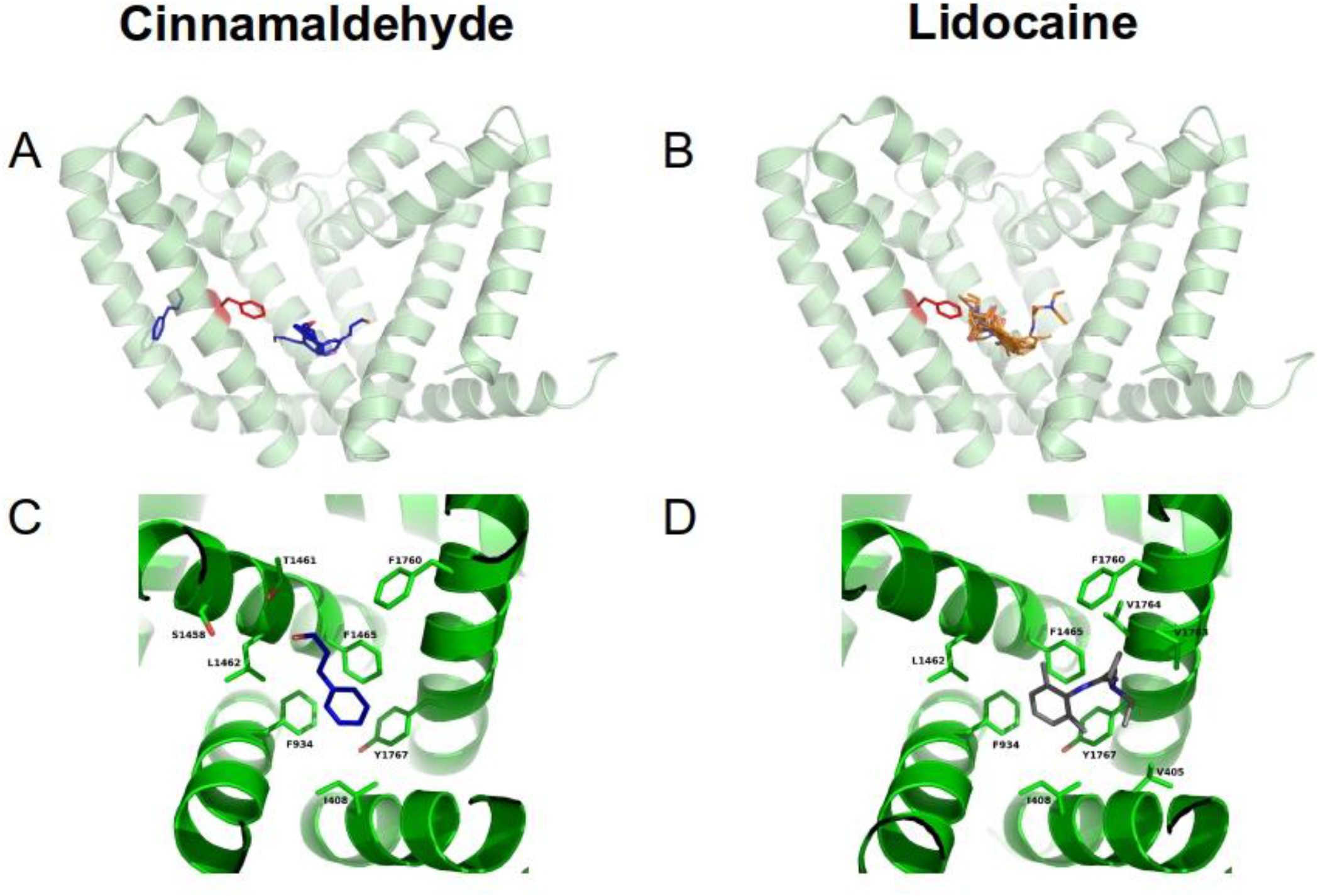
Interacting site of cinnamaldehyde in the hNa_V_1.5 channel. A model with an energetically stable conformation of the hNa_V_1.5 channel was used to run molecular docking simulations. The left panel shows that CA (in blue) was bound at the LA binding site, in close proximity to the residues F934, F1645, F1760 and Y1767. The S6 α-helixes of the four domains of the channel are indicated at the corners of the figure. For comparison, the right panel shows docking simulations of lidocaine (in grey), which was bound to the same pocket.

### 2.3. Comparison of the effect of cinnamaldehyde on the different Na_V_1.5 mutations

Blockade of I_Na_ by CA was reduced in the mutant F1760A. The IC_50_ for I_Na_ blockade was increased to 2.3 ± 0.04 mmol·L^-1^ (H = 1.8 ± 0.06) and was significantly different from the IC_50_ obtained in the WT channel (P < 0.05; Figure 5A). TB with CA 1 mmol·L^-1^ was not significantly different from that obtained in the WT and an increased frequency of stimulation did not further enhance the total block of I_Na_ by CA. (n = 6; Figure 6A and B). Decreasing the HP to -80 mV hardly increased blockade of I_Na_ by 1 mmol·L^-1^ CA (Figure 6C). In control condition in the F1760A mutant, the V_inac_ and V_act_ (and corresponding slope factors) were -70.0 ± 0.2 mV (6.1 ± 0.4 mV) and -43.9 ± 3.8 mV (6.8 ± 0.2 mV), respectively. CA changed neither the V_inac_ (-73.1 ± 0.2 mV) nor the s_inac_ (6.1 ± 0.2 mV; Figure 6D). V_act_ (-44.8 ± 3.9 mV) and s_act_ (5.9 ± 0.5 mV) were also not affected by CA 1 mmol·L^-1^.

**Figure 5.**
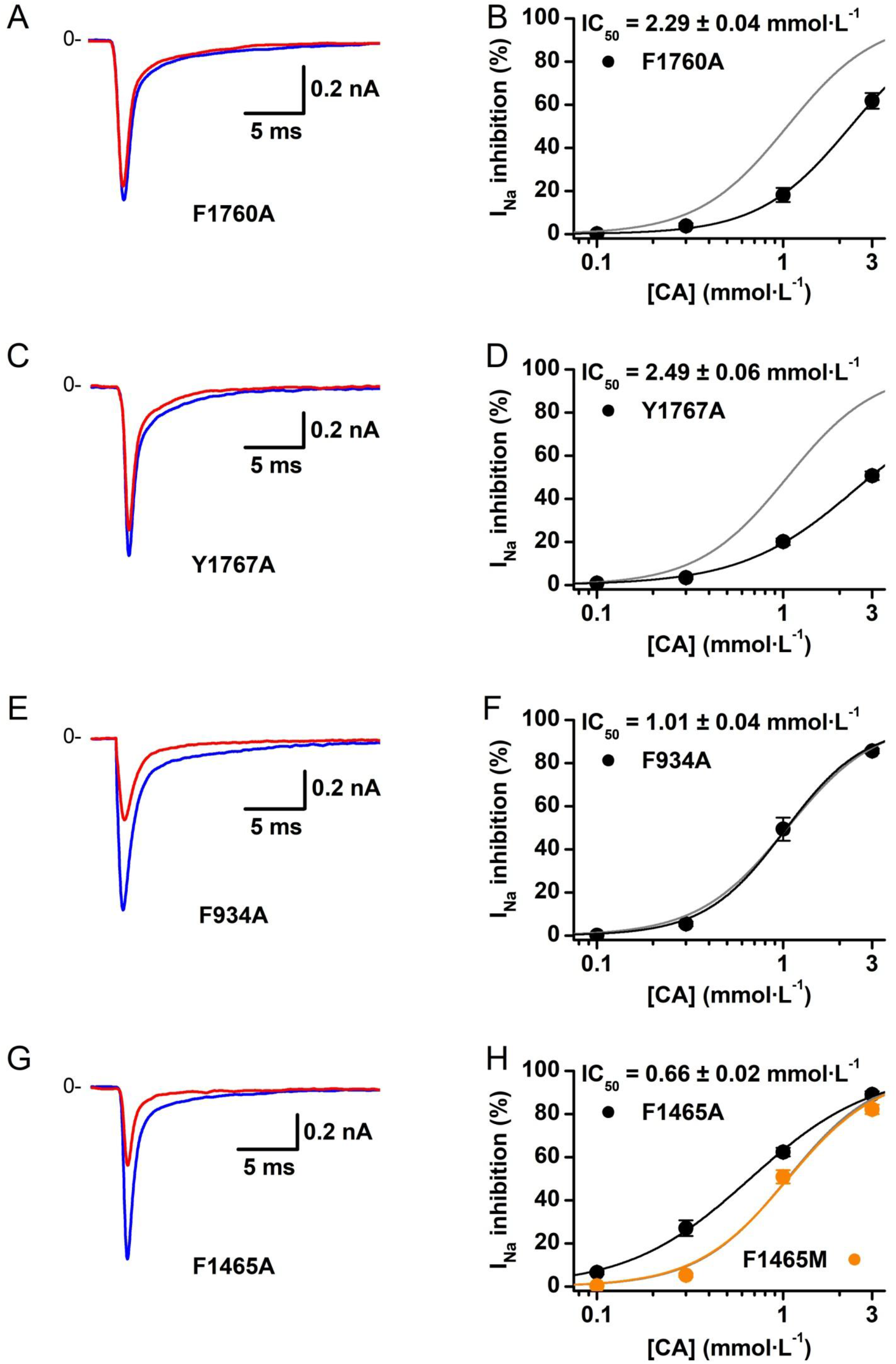
**Effects of several mutations in the LA binding pocket on cinnamaldehyde** blockade of the hNa_V_1.5 channel. A, C, E and G.- representative currents traces showing the effects of CA 1 mmol·L^-1^ on the I_Na_ in the mutants F1760A, Y1767A, F934A and F1465A/M, respectively. Control condition is represented in blue traces and CA 1 mmol·L^-1^ in red. I_Na_ was evoked using 50-ms voltage-clamp pulses from -100 to -20 mV. B, D, F and H.- Concentration-effect relationships of the CA effect on I_Na_ in these mutants. The dots represent the mean ± s.e.m. inhibition percentage with the application of different concentrations of CA. The black lines represent the Hill fit of the data. In each graph the Hill fit obtained in the WT channel (grey lines) is shown for comparison. The respective IC_50_ values are shown in the insets. Note that the blockade of I_Na_ by CA is significantly decreased in the mutations F1760A and Y1767A and, on the contrary, increased in the mutation F1465A (P < 0.05; n = 13; one-way ANOVA with Tukey’s post hoc test). Blockade of hNa_V_1.5 by CA in the mutants F934A and F1465M (in orange in panel H) is similar to WT.

**Figure 6.**
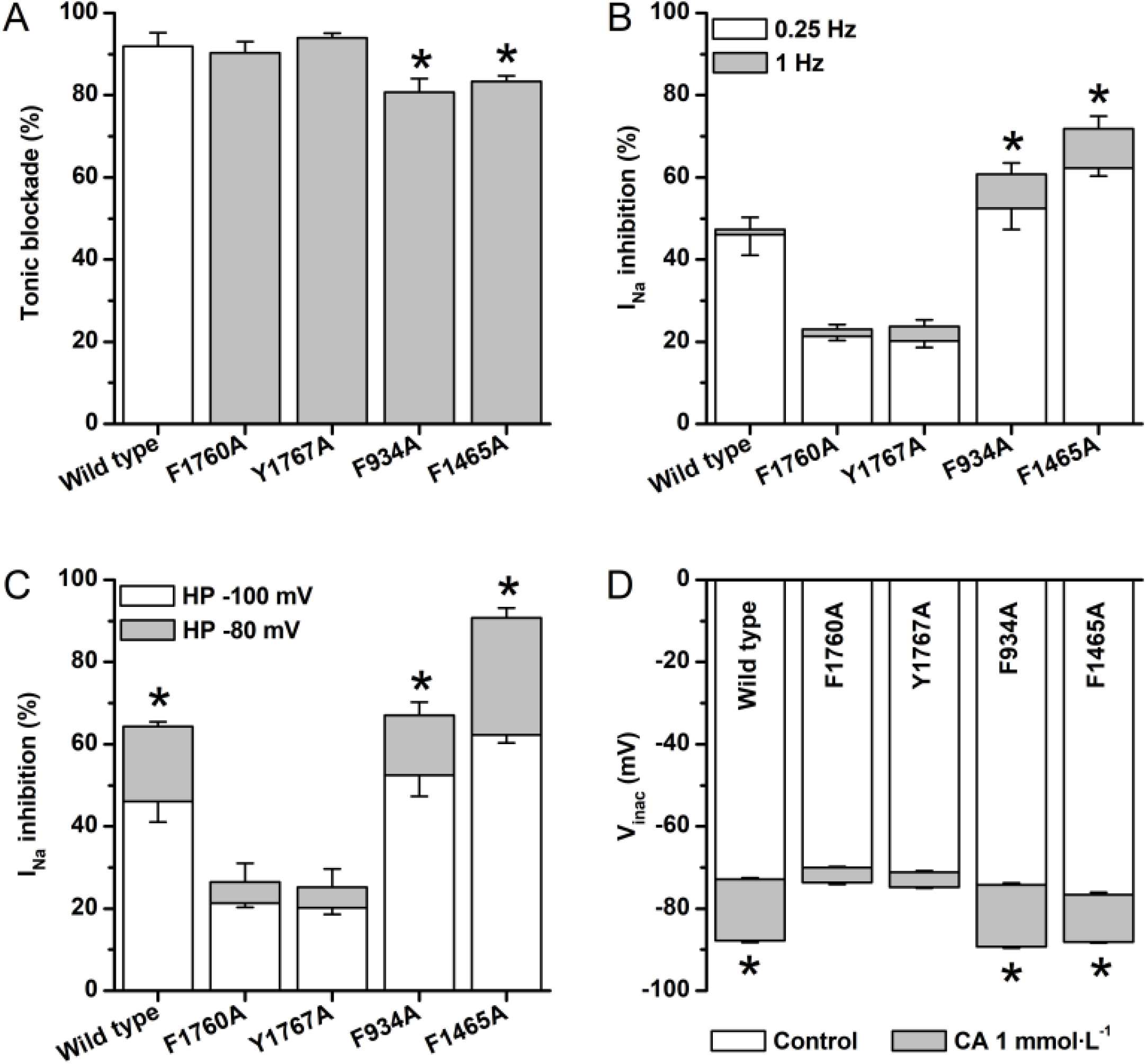
Tonic and voltage-dependent effects of cinnamaldehyde on mutated hNa_V_1.5 channels. In all cases, and for a better comparison, the data of the WT is depicted in dashed lines. A.- Bar graph showing resting block of I_Na_ by CA (1 mmol·L^-1^) in mutants F1760A, Y1767A, F934A and F1465A. Note that resting block was not affected in mutants F1760 and Y1767 but was decreased in mutants F934 and F1465. * P < 0.05 with respect to WT; n = 6; one-way ANOVA with Tukey’s post hoc test. B.- Bar graph illustrating blockade of I_Na_ by CA 1 mmol·L^-^1 at different stimulation frequencies. In white, data obtained at a stimulation frequency of 0.25 Hz and in grey data obtained at a stimulation frequency of 1 Hz. Note that in mutants F934 and F1465 CA increased its blockade at a high stimulation-frequency. * P < 0.05 with respect to its own control (0.25 Hz). § P < 0.05 with respect to WT at 1 Hz; n = 6, two-way ANOVA with Tukey’s post hoc test. C.- Bar graph presenting the effects of decreasing the HP on the blockade of I_Na_ by CA (1 mmol·L^-1^). I_Na_ was evoked using 50-ms voltage-clamp pulses to -20 mV from a HP of -100 mV (in white) and then to the same test pulse but from a HP of -80 mV (in grey). Note that while in mutants F1760A and Y1767A the blockade of I_Na_ by CA was not increased at a HP of -80 mV, however, in the mutants F934 and F1465A the blockade was increased. * P < 0.05 with respect to its own control (HP -100 mV). § P < 0.05 with respect to WT at HP -80 mV; n = 5, two-way ANOVA with Tukey’s post hoc test. D.- Bar graph summarizing the effects of CA on V_inac_. In white, data obtained in control condition and in grey data obtained in the presence of CA 1 mmol·L^-1^. CA induced a shift in the availability curves toward more hyperpolarized potentials. As a result, a ΔV_inac_ is seen as a grey area in the bars. Note that ΔV_inac_ was not significantly increased in mutants F1760 and Y1767 but it was increased in mutants F934 and F1465. * P < 0.05 with respect to its own control condition. § P < 0.05 with respect to WT in CA condition; n = 8, two-way ANOVA with Tukey’s post hoc test. All data is presented as mean ± s.e.m.

For comparison, we studied the effects of lidocaine (LID) on the hNa_V_1.5 channel. LID blocked the WT I_Na_ with an IC_50_ of 150 ± 10 µmol·L^-1^ (H = 0.9 ± 0.07). The IC_50_ for LID blockade of I_Na_ in the F1760A mutant was significantly increased to 3.3 ± 0.3 mmol·L^-1^ (H = ± 0.1; P < 0.05; Figure S1A). To rule out that some of the obtained results could be the consequence of changes in hNa_V_1.5 channel function due to the point mutation, we performed experiments with CdCl_2_, a well-known Na^+^ channel blocker that binds the P-loop site [16]. The obtained IC_50_ for I_Na_ inhibition by CdCl_2_ were 0.19 ± 0.02 mmol·L^-1^ (n = 5) for the WT and 0.21 ± 0.01 mmol·L^-1^ (n = 5) for the F1760A (corresponding H of 1.0 ± 0.1 and ± 0.06). The differences were not statistically significant (Figure S1B).

The present experiments show that the mutation Y1767A significantly increased the IC_50_ of CA blockade of I_Na_ to 2.5 ± 0.06 mmol·L^-1^ (H = 1.4 ± 0.06; P < 0.05; Figure 5C and D). In the Y1767A mutant, I_Na_ blockade was essentially tonic and not frequency dependent as in the WT. However, no significant changes in V_inac_ and s_inac_ (8.4 ± 0.5 vs. 7.4 ± 0.5) were observed in this mutant with CA 1 mmol·L^-1^ and the decrease in HP to -80 mV scarcely increased blockade, as opposed to WT (P < 0.05; Figure 6A – D). Activation parameters V_act_ and s_act_ remained unchanged upon CA application (-35.2 ± 0.4 mV vs. -36.0 ± 0.5 mV and 6.0 ± 0.5 mV vs. 6.5 ± 0.3 mV, respectively). The IC_50_ for LID blockade was significantly increased in the mutant Y1767A (400 ± 31 µmol·L^-1^ with H = 0.8 ± 0.05; P < 0.05; Figure 5.S1A). This, however, was not the case for the IC_50_ for CdCl_2_ blockade of I_Na_ (0.22 ± 0.06 mmol·L^-1^; H = 0.9 ± 0.02; Figure S1B). The mutation F934A, on the other hand, did not change the IC_50_ for CA blockade of I_Na_ (1.01 ± 0.04 mmol·L-1; H = 2.0 ± 0.18; Figure 5.5E and F). Nevertheless, the TB was significantly decreased (P < 0.05; Figure 6A) and the V_inac_ was shifted leftward by ≈ 13 mV (s_inac_ = 7.6 ± 0.4 mV in control and 8.1 ± 0.3 mV in CA; Figure 6D). Consequently, I_Na_ inhibition was enhanced at 1 Hz (P < 0.05; Figure 6B) and at the reduced HP (P < 0.05; Figure 6C). On the contrary, CA (1mmol·L^-1^) did not affect V_act_ and s_act_ (-32.9 ± 1.1 mV vs -34.1 ± 1.2 mV and 5.4 ± 0.9 mV vs 6.2 ± 1.0 mV, respectively).

The IC_50_ for LID inhibition of I_Na_ was not significantly affected in this F934A mutant (149± 15 µmol·L^-1^ with H = 0.6 ± 0.04). Nonetheless, the IC_50_ for CdCl_2_ inhibition of I_Na_ was significantly increased to 0.32 ± 0.03 mmol·L^-1^ (H = 0.9 ± 0.07; P < 0.05; Figure S1B). Surprisingly, I_Na_ blockade by CA was enhanced in mutation F1465A with respect to WT. The IC50 was decreased to 0.66 ± 0.02 mmol·L^-1^ (H = 1.3 ± 0.05; P < 0.05; Figure 5G and H), together with a decrease in TB and, as a consequence, a slight increase in the UDB (P < 0.05; Figure 6A and B). CA (1 mmol·L^-1^) exerted a shift of V_0.5_ for I_Na_ availability of ≈ 12 mV in the hyperpolarizing direction (s_inac_ = 7.1 ± 0.5 mV in control and 7.7 ± 0.3 mV in CA conditions) and a significantly increased blockade at a reduced HP of -80 mV with respect to WT (P < 0.05; Figure 6C and D). Fitting parameters from the activation curve (V_act_ and s_act_) were not affected by CA 1 mmol·L^-1^ (-43.1 ± 1.0 mV vs -45.4 ± 1.1 mV and 5.4 ± 0.8 mV vs 5.7 ± 0.9 mV, respectively). The F1465A I_Na_ blockade by LID was also enhanced. Its IC_50_ was decreased to 99 ± 13 µmol·L^-1^ (H = 0.7 ± 0.05; P < 0.05; Figure S1A). The decrease proportion in IC_50_ for LID was 0.6, exactly the same to the one obtained for CA. I_Na_ blockade by CdCl_2_ was not affected (IC_50_ = 0.24 ± 0.08 mmol·L^-1^ and H = 1.0 ± 0.02). The decrease in IC_50_ for CA and LID in F1465A mutation prompted us to particularly study the influence of this residue as a steric hindrance for the interactions of CA with other residues of the LA pocket. The use of the longer chain substitute methionine partially rescued the WT phenotype. In the F1465M mutant, the IC_50_ for I_Na_ blockade by CA was 1.05 ± 0.09 mmol·L^-1^ (H = 1.7 ± 0.3; Figure 5H) and the TB, accounted for 91.7 ± 1.6% of the total blockade. The V_inac_ (and s_act_) in control condition was -72.9 ± 0.4 mV (7.6 ± 0.8 mV) and it was shifted by CA (1 mmol·L^-1^) to -83.0 ± 0.7 mV (7.1 ± 0.4 mV). The blockade at more depolarized HP was similar to the WT (65.4 ± 1.6%) as well as the estimated V_act_ and s_act_ were also similar to that of the WT (-34.8 ± 0.7 mV and 6.5 ± 0.6 mV, respectively) and were not modified by CA (V_act_ = -32.6 ± 0.9 mV and s_act_ = 6.0 ± 0.7 mV). I_Na_ blockade by LID was partially recovered to an IC_50_ close to WT (127 ± 17 µmol·L^-1^ with H = 0.8 ± 0.08). I_Na_ blockade by CdCl_2_ was not significantly changed (IC_50_ = 0.16 ± 0.09 mmol·L^-1^ and H = 1.0 ± 0.06; Figure S1B).

### 2.4. Double point mutation F1760A/Y1767A diminishes cinnamaldehyde effects on hNa_V_1.5

Since both F1760 and Y1767 residues seem to be important for CA blockade of hNa_V_1.5, we constructed the double mutant F1760A/Y1767A. The estimated IC_50_ for CA in the double mutant was ≈ 11 mmol·L^-1^, with an I_Na_ blockade at 3 mmol·L^-1^ of less than 20%. The small effect of 1 mmol·L^-1^ CA on I_Na_ (8.5 ± 1.3% inhibition) was essentially tonic, as in the WT, accounting for a 94.6 ± 0.5% of the total block. Increment of the stimulation rate did not enhanced the total block of I_Na_ by CA 1 mmol·L^-1^ (6.7 ± 0.9%). CA blockade was hardly increased by a depolarized HP (8.1 ± 0.7%). Activation and inactivation curves were not changed by CA. Figure 7A - D summarizes the results obtained with the double mutant F1760A/Y1767A. Blockade of I_Na_ by CdCl_2_ was not affected (IC_50_ of 0.18 ± 0.01 mmol·L^-1^ with H = 1.1 ± 0.06; Figure S1A) but the IC_50_ for LID was increased to 3.3 ± 0.2 mmol·L^-1^ (H = 0.8 ± 0.04; Figure S1B).

**Figure 7.**
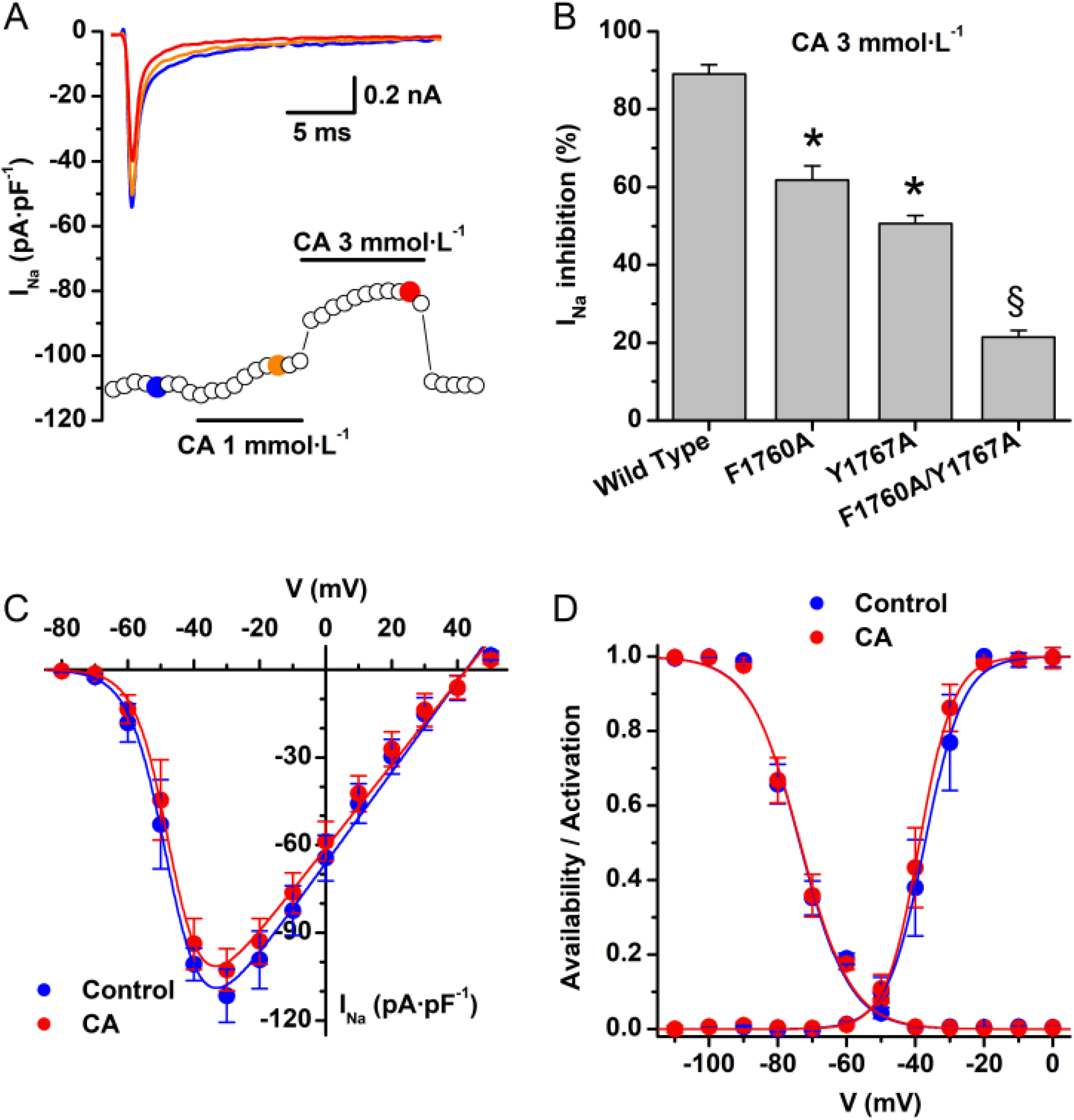
Cinnamaldehyde blockade of I_Na_ is almost abolished in the double mutant. F1760A/Y1767A channel. A.- Time course of the CA effect on the I_Na_ from a HEK293T cell expressing the double mutant F1760A/Y1767A hNa_V_1.5 channel. CA 1 and 3 mmol·L^-1^ was applied as indicated. Representative current traces in each condition are shown in the inset and correspond with the coloured data points (control in blue and CA 1 and 3 mmol·L^-1^ in orange and light red, respectively). For clarity only the first 25 ms of the current traces are shown. B.- Bar graph depicting the blockade of I_Na_ by the maximal concentration of CA. The effects of CA 3 mmol·L^-1^ were markedly diminished in the double point mutation F1760A/Y1767A. Data is presented as mean ± s.e.m.* P < 0.05 with respect to WT. § P < 0.05 with respect to all other conditions, including WT; n = 13, one-way ANOVA with Tukey’s post hoc test. C.- Current-to-voltage relationships of I_Na_ in the double mutant under control condition (blue) and in the presence of CA 1 mmol·L^-1^ (red). The dots represent the mean ± s.e.m. (n = 8). D. Availability and activation curves for F1760A/Y1767A in control (blue) and in the presence of CA 1 mmol·L^-1^ (red). The dots represent the mean ± s.e.m. (n = 8). The V_inac_ and the s_inac_ in control and in the presence of CA 1 mmol·L^-1^ were similar (-74.4 ± 0.8 mV vs. -74.3 ± 0.7 mV and 7.4 ± 0.7 mV vs. 7.5 ± 0.6 mV). The V_inac_ and the s_inac_ were also not changed by CA (-37.2 ± 0.4 mV vs. -38.9 ± 1.4 mV and 5.7 ± 0.5 mV vs. 5.0 ± 0.5 mV, respectively).

## 3. Discussion

The main outcome of the present investigation is that cinnamaldehyde exhibited LA like actions on human Na_V_1.5 channels expressed in HEK293T cells. Blockade of I_Na_ was concentration- and voltage-dependent and occurred in an almost tonic fashion. Residues F1760 and Y1767 are important for CA blockade and a double point mutation F1760A/Y1767A abolished the blockade of I_Na_ by CA.

As a TRPA1 agonist [5], CA has been used in pain and neurogenic inflammation research. However, we previously showed that CA exhibits an intriguing blocking action on the bacterial voltage-gated Na^+^ channels NaChBac [6]. In the present experiments, CA blocked hNa_V_1.5 channels, transiently expressed in HEK293T cells. I_Na_ blockade was voltage-dependent and occurred with minor effects on the inactivation kinetics of I_Na_. These effects are reminiscent of those of LID on hNa_V_1.5 channels expressed in HEK293T cells and on TTX-resistant Na^+^ currents in DRG neurons [17, 18]. As for LID [19], blockade of hNa_V_1.5 by CA at 1 Hz stimulation rate, was strongly tonic, not use-dependent. The strong tonic block in our conditions, suggests that CA binds with high affinity to the rested state conformation of the Na^+^ channel at the HP of -100 mV. As for most LA compounds, the blocking action of CA on WT hNa_V_1.5 was voltage-dependent, i.e. block was enhanced by ≈ 15% at 1 mmol·L^-1^ CA at depolarized holding potentials (-80 mV). This was consistent with a shift of ≈ 12 mV to more hyperpolarized potentials of the V_inac_, an action that is typical of LA, in particular LID [8, 14]; the V_act_, however, was not shifted by CA. A plausible explanation for our findings is that CA interacts with high affinity with the resting state conformation of the Na^+^ channel that follows voltage-dependent transitions to an absorbing inactivated state [20]. Our docking simulation on a structural model of the hNa_V_1.5 channel, indicate that both LID and CA bind in the vicinity of the F1760 residue in the transmembrane segment S6 of the domain IV. This residue has been identified as the main structural determinant of the high-affinity block by local anaesthetics [8, 9, 21], thus suggesting a common site of action on the hNa_V_1.5 channel.

The results of these simulations were confirmed by patch-clamp experiments where it was shown that the F1760A mutation markedly reduced hNa_V_1.5 channel block by CA, thus providing strong support for a common site of action of CA and LID. Contrary to WT, in F1760A channels the voltage-dependent effects of CA were greatly attenuated: decreasing the holding potential barely increased blockade of I_Na_ by CA. Consequently, the V_inac_ was not significantly shifted. These results suggest that besides its role in the “LA binding pocket”, residue F1760 could be also important for the voltage-dependent effects of CA on the hNa_V_1.5 channel. We may speculate that changing this residue could set a channel conformation that interacts with CA in a less voltage-dependent manner. However, the basic functions in the F1760A mutated hNa_V_1.5 channels were not compromised by the mutation since its inhibition by CdCl_2_ resulted identical to that of the WT with an IC_50_ in the same range as that previously reported [22]. These results confirm the idea [6] that due to their structural similarity, LID and CA share a common interaction site within the hNa_V_1.5 channel (the residue F1760), but they also provide evidence that other residues could also interact with CA. Although the residue F934 is close to the CA binding site, it is not relevant to CA action. The F934A mutant behaved essentially as the WT channel with the exception that the UDB of I_Na_ by CA was relatively increased. This effect could be due to changes in the properties of the channel due to the mutation as the IC_50_ for CdCl_2_ blockade of I_Na_ was significantly increased in this mutant. On the other hand, the mutation F1465A increased the potency of CA on the hNa_V_1.5 current. In this mutation the UDB was also increased as well as the blockade of I_Na_ at higher stimulation frequency. These effects may be partially responsible for the higher sensitivity of this mutant to CA. Interestingly, in this mutation, the decrease in the IC_50_ for LID and for CA occurred in the same proportion (0.6 fold). We may hypothesize, then, that the large phenylalanine in position 1465 could be a steric hindrance that would obstruct the access of CA to the LA pocket. When the smaller alanine is present, the access of CA to the other residues is eased and I_Na_ block by CA is stronger. The longer chain methionine at this position seems to confirm this hypothesis since in the F1645M mutation, CA potency on I_Na_ is similar to WT. As previously shown for other LA-like drugs [15, 23], the residue Y1767 is directly involved in blockade of hNa_V_1.5 by CA. Alanine mutation in this residue increased the IC_50_ for CA blockade by approximately 2.5-fold and the small effect was not voltage-dependent with almost no change in the V_inac_. As with the F1760A mutation, TB was not changed. That the residues F1760 and Y1767 are directly involved in CA block was confirmed with the double point mutation F1760A/Y1767A. In this double mutant the IC_50_ for CA blockade was increased about 10-fold, almost totally abolishing the effect of CA. Again, the small action of CA was not voltage-dependent and TB block was not changed. The IC_50_ for LID blockade on F1760A/Y1767A was increased more than 20-fold. Interestingly, the Hill fit of the concentration-effect relationship for the inhibitory action of LID on the double mutant overlaps with the one obtained on the single point mutation F1760A. This may suggest that for the action of LID only the F1760 residue is important, whereas for CA both residues (F1760 and Y1767) are taking part in the interaction with this compound. Residues in DIV that contribute to the LA receptor site face the inner pore region of the channel [8]. They may become accessible to the LA molecules as a result of the conformational changes and stabilize drug binding by interactions with charged, hydrophobic, and/or aromatic residues on the drug molecules [21, 24-26]. It is possible then that differences in efficacy of CA and other LA molecules are a consequence of restrictions to receptor access.

The concentrations of CA used in the present study can be considered high. However, we must note that cinnamon extracts, whose main component (∼90%) is CA, are commonly used as flavorants at concentrations ≥ 0.5 % (several millimoles per liter; [27]. Cinnamon extracts have been also used in clinical trials to lower blood glucose levels in diabetic individuals [28-32] at concentrations that could be equivalent to those used in the present experiments. It is therefore important to uncover the possible undesirable side-effects of CA that are relevant to human health.

In conclusion, our findings strongly indicate that CA and LID share common structural determinants for the inhibition of Na^+^ channels and further support our conclusion that CA has LA-like actions. Our study should be interpreted as a thorough characterization that unveils an unrecognised pharmacological property of CA as a Na_V_1.5 blocker with LA-like properties.

## 4. Materials and Methods

### 4.1. Human embryonic kidney cells culture and transfection

Human embryonic kidney cells, HEK293T, were seeded on 18 mm glass coverslips previously coated with poly-L-lysine (0.1 mg·L^-1^). Cell media consisted in a Dulbecco’s modified Eagle’s medium (GIBCO, NY, USA) containing 10% of human serum, 2 U·mL^-1^ penicillin, 2 mg·mL^-1^ streptomycin and 2 mmol·L^-^1 L-glutamine. The cells were stored at 37 °C in a humidity-controlled incubator with 10% CO_2_. Na_V_1.5 mutations were introduced by overlap extension polymerase chain reaction (PCR) and the amplicons containing the mutation were cloned into the pCAGGS-IRES-GFP vector [17] using appropriate restriction sites and sequence-verified. The genes encoding human Na_V_1.5 wild type (WT) or the mutants R1623Q, F934A, F1465A, F1465M, F1760A and Y1767A as well as the double mutant F1760A/Y1767A were transiently transfected in HEK293T cells using the TransIT-293 reagent (Mirus, MI, USA). We expressed only the pore-forming α subunit of the Na_V_1.5 channel because this suffices to generate I_Na_ similar to the native ones [19] and the mutations studied occur in this subunit. Additionally, this approach is commonly reported in the literature and allows us to compare with other results. Depending on the mutation, only 0.1 µg – 0.3 µg of Na_V_1.5 cDNA were used for transfection.

### 4.2. Patch-clamp recordings in HEK293T cells

Coverslips with seeded cells were placed in a recording chamber on the stage of an inverted microscope, with an Ag|AgCl wire as the reference electrode. Extracellular solutions were perfused by gravity via a multi-barrelled pipette. Transfected cells were identified by the GFP expression during the patch-clamp experiments, performed 24 - 48 h after transfection. Longer culture periods resulted in huge Na^+^ currents making difficult to control membrane potential in patch-clamp experiments. The cells were let to stabilize for 2 minutes, before beginning the recordings, in a Krebs solution containing (in mmol·L^-1^): 150 NaCl, 6 KCl, 1 MgCl_2_, 1.5 CaCl_2_, 10 glucose, 10 HEPES and titrated to pH 7.4 with NaOH. Whole-cell currents were recorded in only one cell per coverslip. Measurements were performed at room temperature using an EPC7 patch-clamp amplifier (LIST Electronics, Darmstadt, Germany), a TL-1 DMA interface (Axon Instruments) and the pClamp software (Version 9.0, Axon Instruments, CA, USA). Currents were filtered at 3 kHz, digitized at 50 μs intervals, stored on a computer and analysed off-line with the WinASCD software (KU Leuven, Belgium). The extracellular solution for HEK293T cells contained (in mmol·L^-1^): 140 NaCl, 10 HEPES, 2 CaCl_2_, 1 MgCl_2_, and 10 glucose, pH 7.4 with NaOH. The pipette solution contained (in mmol·L^-1^): 130 CsCl, 5 Na_2_ATP, 5 Na_2_-creatine phosphate, 5 EGTA, 1 MgCl_2_, 1 CaCl_2_ (free Ca^2+^, ≈ 0.4 nmol·L^-1^) and 10 HEPES, pH adjusted to 7.2 with CsOH. Patch pipettes (1.2 - 1.5 MΩ) were pulled from borosilicate capillary tubes. Rs (3.6 ± 0.5 MΩ) and Cm (21.7 ± 0.6 pF) could be determined by the usual routines and the built-in compensation circuits of the EPC-7 amplifier. In all recordings Rs was electronically compensated up to 50% without ringing and was continually monitored during the experiment. Liquid junction potential was compensated before establishing the gigaseal and capacity transients were cancelled using the pClamp P/4 protocol. I_Na_ was monitored using 50-ms voltage pulses to -20 mV, applied at 0.25 Hz from a HP of -100 mV. A more physiological HP of -100 mV was chosen because peak I_Na_ densities were similar at HP of -120 mV and -100. Current amplitudes of each cell were normalized to the Cm and expressed as current densities (pA·pF^-1^). Due to the presence of a residual Rs, an adequate characterization of the fast I_Na_ activation is not possible. Thus, we only estimated the time-to-peak of the current.

### 4.3. Patch-clamp protocols and data analysis

Peak I_Na_ amplitude was measured as the difference between peak inward current and the zero-current level. The concentration-effect relationships for the effects of CA on I_Na_ were fitted by a Hill function of the form:

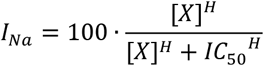

where [x] is the concentration of the compound used, IC_50_ is the effective inhibitory concentration and H is the Hill coefficient. Current-voltage relationships (I-V) and availability curves were obtained by clamping the cells at a HP of -100 mV and using a standard double-pulse voltage-clamp protocol (frequency, 0.25 Hz). From the HP, a 50 ms test pulse to -20 mV was preceded by 500 ms prepulses to various membrane potentials. The time interval between pulses was 1 ms. Availability curves were obtained from the normalization of the amplitude of currents recorded at the test pulse by the maximal current (I_Max_). These curves were fit by the equation:

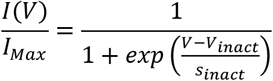

where V_inac_ is the voltage for half-maximal availability and s_inac_ is the slope factor. I-V relationships were obtained from the amplitude of currents elicited by pre-pulse potentials and were fit by the equation:

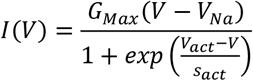

where G_Max_ is the maximal whole-cell conductance, V_Na_ is the equilibrium potential for Na^+^, V_act_ is the voltage for half-maximal activation and sact is the slope factor. Activation curves were obtained for each cell by dividing the experimental current amplitudes, I(V), by the corresponding values of G_Max_(V – V_Na_). Activation curves were fit by the equation:

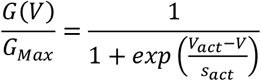

At the test potential, the time course of the inactivation phase of the I_Na_ traces were fitted by a double exponential:

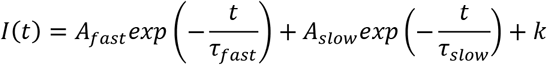

This procedure yielded two inactivation time constants, τ_fast_ and τ_slow_ and the amplitudes of the two components (A_fast_ and A_slow_), from which we calculated the relative amplitude of the slow component. k is a constant that represents the steady-state current amplitude. This value was, however, not discussed because it was negligible (< -6·10^-6^ pA·pF^-1^).

In order to investigate tonic (TB) and (UDB) use-dependent blockades of I_Na_ by CA, cells were voltage-clamped with 50-ms pulses from -100 mV to -20 mV at a frequency of 0.25 Hz. Stimulation was stopped and perfusion with CA was then initiated. After 1 minute, stimulation was resumed (in the presence of CA) at frequencies of 0.25 or 1 Hz and then switched to 0.25 Hz. TB was estimated as the difference between peak control I_Na_ and the I_Na_ at the first pulse after resumption of stimulation during drug superfusion. UDB was considered to be the difference between peak I_Na_ at the first and the 15th pulse after drug exposure. TB and UDB are expressed as percentage of total block which was estimated as the difference between peak I_Na_ recorded in control and CA-containing condition.

### 4.4. Molecular modelling and docking simulations

We used the software Modeller (University of California, USA)^208^ to generate a model of the structure of hNa_V_1.5 using its bacterial homologue Na_V_AB (PDB code: 3RVY) [33, 34] as template. This model was further refined, using NAMD (University of Illinois, USA) [35], by running 10 ns molecular dynamics simulations to obtain an energetically stable conformation. The final model is composed by the four repeats of the segments of S5, pore forming and S6 (S5-pore forming-S6), with repeat I comprising the residues from 231 to 415, repeat II 820 to 941, repeat III 1315 to 1472 and repeat IV 1638 to 1774. This model was then used as receptor for in silico docking studies to analyze its interaction with CA using the program AutoDock Vina (University of California, USA) [36]. The complete pore was used as search space allowing flexibility to the side chain pointing inwards as well as for CA.

### 4.5. Chemicals

All chemicals were purchased from Sigma-Aldrich (Bornem, Belgium). For CA nd lidocaine, we took advantage that both CA and LID are soluble in water at ∼ 10 and 3 mmol.L^-1^, respectively (https://go.drugbank.com/), and dilutions of these chemicals were prepared daily in extracellular bath solution from the stocks that were kept at 4° C.

### 4.6. Statistical analysis

Data was analysed, and fitted, using the routines of Origin 9.0 (OriginLab Corporation, MA, USA) and expressed as means and standard errors of means (mean ± s.e.m.) from n cells. Group data subjected to analysis had n values ≥ 5 independent samples per group and compared groups had equal size. Normality was confirmed with the ShapiroWilk test (95% confidence level). Statistical significance was evaluated with a paired sample t-test within groups or two-sample t-test between groups. For differences between multiple groups of data we used a one-way or two-way ANOVA, according to the experimental situation, followed by a Tukey’s post hoc test. If conditions for parametric tests were not met, non-parametric tests were used. Tests used for statistical comparison between groups are indicated either in the text or in the figure legends. Differences were considered statistically significant if P < 0.05.

## Author Contributions

Conceptualization, A.T. and K.T.; methodology, K.T.; software, A.T.; validation, K.T. and J.L.A.; formal analysis, J.A.C., A.T. and J.L.A.; investigation, J.A.C., A.L.R. and A.T.; resources, K.T.; data curation, J.A.C., A.T. and J.L.A.; writing—original draft preparation, J.A.C., J.L.A., K.T.; writing—review and editing, J.L.A., K.T.; visualization, J.A.C., J.L.A.; supervision, K.T.; project administration, K.T.; funding acquisition, K.T. All authors have read and agreed to the published version of the manuscript.

## Institutional Review Board Statement

Not applicable.

## Informed Consent Statement

Not applicable.

## Data Availability Statement

The original contributions presented in this study are included in the article/Supplementary Material. Further inquiries can be directed to the corresponding authors.

### Acknowledgments

The authors wish to thank Prof. Angelika Lampert (Institute of Physiology, RWTH Aachen University) for kindly providing us the hNa_V_1.5 clone and Melissa Benoit for excellent assistance and support.

---

**Figure S1.**
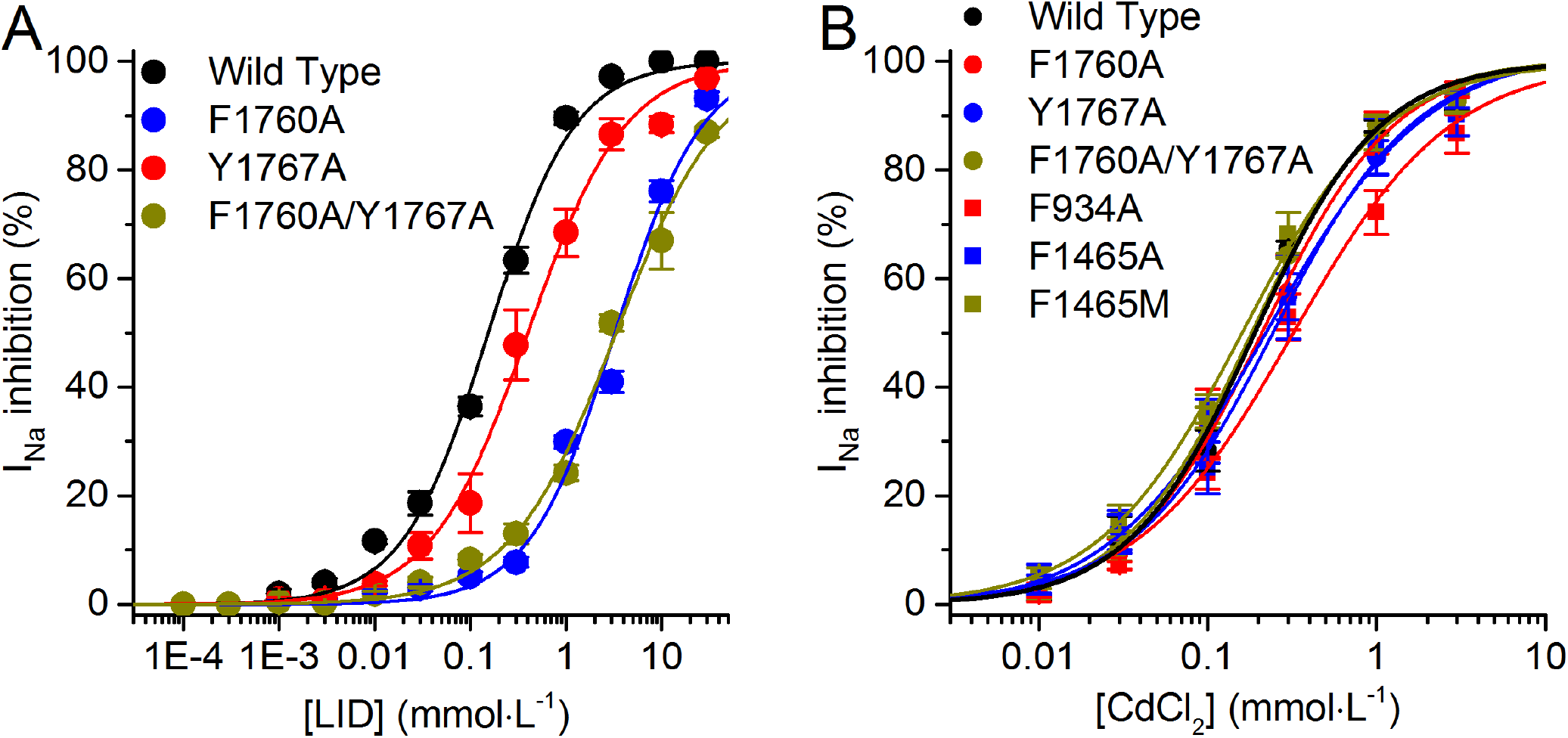
Concentration dependent effects of lidocaine and CdCl_2_ on the currents carried by WT and several channel mutants.

